# Triterpenoid production with a minimally engineered *Saccharomyces cerevisiae* chassis

**DOI:** 10.1101/2022.07.11.499565

**Authors:** Hao Guo, Simo Abdessamad Baallal Jacobsen, Kerstin Walter, Anna Lewandowski, Eik Czarnotta, Christoph Knuf, Thomas Polakowski, Jérôme Maury, Christine Lang, Jochen Förster, Lars M. Blank, Birgitta E. Ebert

**Affiliations:** Australian Institute for Bioengineering and Nanotechnology, The University of Queensland, St Lucia QLD, Australia; Institute of Applied Microbiology (iAMB), Aachen Biology and Biotechnology (ABBt), RWTH Aachen University, Aachen, Germany; Novo Nordisk Foundation Center for Biosustainability, Danish Technical University, Lyngby, Denmark; Organobalance GmbH, Berlin, Germany

## Abstract

Triterpenoids, one of the most diverse classes of natural products, have been used for centuries as active ingredients in essential oils and Chinese medicines and are of interest for many industrial applications ranging from low-calorie sweeteners to cosmetic ingredients and vaccine adjuvants. However, not only can the extraction from plant material be cumbersome due to low concentrations of the specific triterpenoid, but concerns are also increasing regarding the sustainability of wild plant harvest while meeting market demands. The alternative is to produce triterpenoids with engineered microbes. Here, we present a generally applicable strategy for triterpenoid production in the yeast *Saccharomyces cerevisiae* based on a modified oxidosqualene cyclase Erg7. The modification reduces the flux into the sterol pathway while increasing the precursor supply for triterpenoid production. The minimally engineered strain was exploited for the exemplary production of the lupane triterpenoids betulin, betulin aldehyde, and betulinic acid at a total titer above 6 g/L, the highest reported so far. To further highlight the chassis concept, squalene, oleanane- and dammarane-type triterpenoids were synthesized to titers at a similar gram scale. We propose the developed baker’s yeast as a host for the thousands of triterpenoid synthesis pathways from plants, reducing the pressure on the natural resources.

## Introduction

Natural products form the biologically relevant subset of the chemical space, which emerged through natural evolution for a myriad of applications, covering such different properties as sun protection, water binding, and pathogen resistance. They represent an invaluable source of bioactive, small molecules for pharmaceutical development and, with insights into their natural biological role, enable more streamlined lead identification than screening vast combinatorial chemical libraries. Pharma, cosmetic, and food industry, besides others, take advantage of these properties provided by functionally tested natural products. One such class of natural products is triterpenoids, widely found in plants but also present in mushrooms and sea cucumbers.^2, 3^ For example, the triterpenoid betulin is a European registered molecule for wound healing and is an active ingredient and emulsifier in ointments for the treatment of challenged skin.^4^ Many other triterpenoids are investigated for a plethora of applications, including betulinic acid and lupeol.^5, 6^

Triterpenoids are the largest natural product class, with more than 20,000 molecules identified to date, derived from a cascade of reactions that leads to a combinatorial explosion. Briefly, after the cyclization of 2,3-oxidosqualene or squalene to mono-up to pentacyclic triterpenoids, these molecules are modified by cytochrome P450 monooxygenases (CYPs), glycosyl- and acyltransferases in cascades of more than ten enzymatic steps. Since the diversity of oxidosqualene cyclases, CYPs, and glycosyltransferases can be combined almost freely, and some cyclases are multifunctional, an immense and structural diverse biochemical space can be synthesized.^7, 8^ This vast diversity variety of triterpenoids might be explained by a biochemical arms race between the host and its respective pathogens.^9^

A challenge of studying natural products is accessibility because their concentrations in natural sources are often insufficient, and they are present in a complex mixture of very similar molecules.^10^ For example, in the various species of the plant ginseng, central in traditional Chinese medicine, about 150 triterpenoids, ginsenosides, have been detected. But even with this very prominent medicinal plant, the molecules with the greatest efficacy are largely unknown, as dedicated studies with single triterpenoids in humans are largely missing.^11^ Again, the availability of triterpenoids restricts their further characterization and consequently their use – a classic catch-22. Yeast, especially *Saccharomyces cerevisiae*, has proven to be an advantageous host for the production of triterpenoids, partly because this Eukaryote provides the triterpenoid precursor 2,3-oxidosqualene via the endogenous mevalonate pathway.^12^ While many plant metabolic pathways have been elucidated and new-to-nature triterpenoids have been synthesized through combinatorial biosynthesis, the amount produced often remained in the low milligram-per-liter range.^13^ In the last couple of years, a single exception was reported, with a titer of about 10 g/L.^14^ The massive genome editing required included the deregulation and overexpression of most genes of the mevalonate pathway.^14^ Most metabolic engineering designs are inspired by work on artemisinic acid production, a sesquiterpene that was industrially produced for some time, with reported titers of above 40 g/L. Artemisinic acid clearly benefitted from the fact that sesquiterpenes are generally secreted into the growth medium, while most triterpenoids are not.^15^

We argue that a yeast strain with minimal genomic manipulations yet effective for a high flux through the mevalonate pathway towards 2,3-oxidosqualene is the ideal chassis for triterpenoid synthesis. With such a chassis in hand, the researchers can focus entirely on the triterpenoid-specific genes and their genetic implementation and rapidly produce quantities that enable further characterizations and initial application testing. Here, we present such chassis that carries a novel variant of the lanosterol synthase Erg7, which results in reduced flux into sterol biosynthesis. In combination with ethanol as carbon source, this chassis strongly accumulated 2,3-oxidosqualene, the precursor of plant-derived triterpenoids such as the lupane-, oleanane-, and dammarane-type triterpenoids studied here. These triterpenoids were produced at summed concentrations above 5 g/L, allowing easy purification of amounts suited for further characterizations. The presented yeast-based triterpenoid platform will improve the accessibility of these complex molecules and thus accelerate the exploitation of this fascinating class of natural products.

## Results

### Downregulation of lanosterol synthase activity is critical for 2,3-oxidosqualene accumulation

A sufficient supply of the central precursor 2,3-oxidosqualene is a prerequisite for producing any triterpenoid in significant quantities. To overproduce this metabolite in *S. cerevisiae*, we deregulated the native mevalonate pathway by overexpressing a truncated version of the 3-hydroxy-3-methyl-glutaryl-coenzyme A (HMG-CoA) reductase Hmg1, a known, rate-limiting step of the pathway (Fig. 1).^16^ This cytosolic enzyme variant lacks the membrane-binding region and is no longer subject to native protein degradation. t*HMG1* overexpression alone resulted in intracellular squalene accumulation, as reported previously, but 2,3-oxidosqualene was not detected in the cell extracts (strain SQ1; data not shown).^16^ To enforce 2,3-oxidosqualene accumulation, we aimed to destabilize lanosterol synthase Erg7, drastically reducing the flux into the native ergosterol synthesis pathway downstream of the intermediate 2,3-oxidosqualene and hence its conversion. Two strategies were tested to control lanosterol synthase activity, which targeted protein degradation or gene expression.

**Figure 1.**
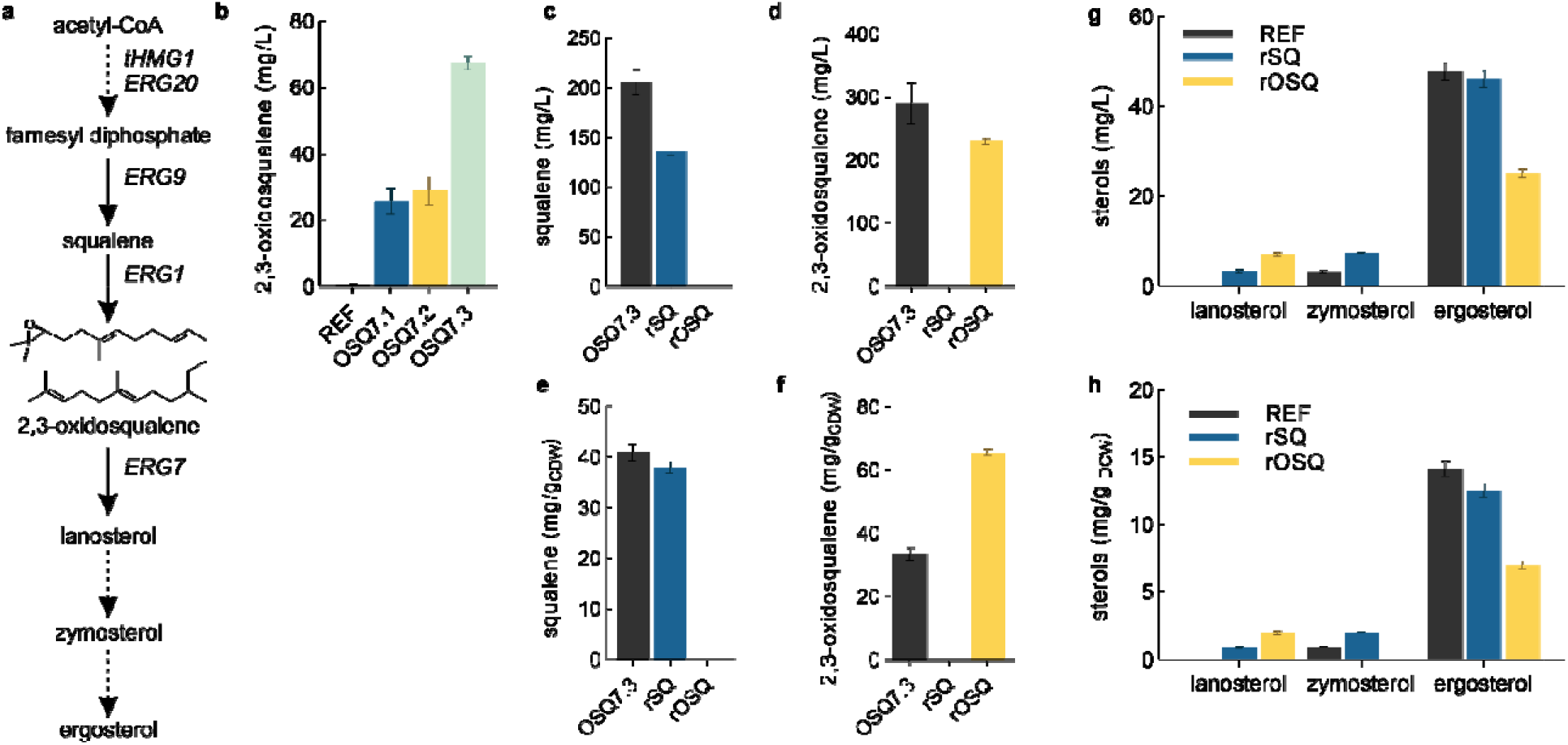
2,3-oxidosqualene biosynthesis via mevalonate and ergosterol pathway **(a)**, volumetric 2,3-oxidosqualene concentrations in the reference *S. cerevisiae* CEN.PK 102-5B (REF), and three clones of this strain with engineered lanosterol synthase, Erg7, depletion **(b)**, volumetric and specific 2,3-oxidosqualene concentrations of mutant OSQ7.3 and the reversed engineered strain rOSQ and its reference rSQ **(c-f)**, volumetric and specific sterol concentrations in REF, rSQ and rOSQ **(g-h)**.

We altered the transcriptional control of *ERG7* by exchanging the native *ERG7* promoter for the promoter of the *HXT1* gene, encoding a low-affinity glucose transporter. Previous studies have used this promoter to redirect metabolism into a heterologous pathway.^17^ However, in an initial evaluation, the resulting strain OSQ11 showed growth only during the glucose phase, in which the *HXT1* promoter ensures sufficient expression of lanosterol synthase. When glucose depleted, gene expression was suppressed to such an extent that cell growth ceased, however, without accumulation of OSQ (data not shown).

The protein depletion strategy used the Tabacco Etch Virus (TEV) protease-induced protein instability (TIPI).^18^ In this approach, protein degradation is achieved by tagging the target protein with a degron cassette containing an internal TEV protease site and dormant unstructured degron elements. In the uninduced state, the target enzyme is functional and present at natural concentrations within the cell. Induction of TEV protease expression leads to targeted proteolysis and consequent uncovering of the degron elements, resulting in rapid degradation of the protein. This strategy ensures a rapid switch from the biomass production phase to the triterpene production phase. The TIPI degron cassette was cloned directly upstream of the *ERG7* stop codon in strain SQ1, resulting in strain OSQ6. The copper-inducible promoter p_*CUP1*_ was cloned upstream of the TEV protease gene into the integrative USER vector pCFB258, and this construct was integrated into OSQ6, resulting in strain OSQ7. OSQ7 was cultured in mineral salt medium, and induction of TEV protease expression was tested with varying copper concentrations. As already suggested from the promoter screening experiments (Supporting Information), the TEV protease appears to be very active even in the non-induced state (0 mM), accumulating 2,3-oxidosqualen e at all concentrations tested (Supporting Information). In addition, there was wide variability among the transformants tested (Fig. 1b), with clone OSQ7.3 having more than twice the OSQ accumulation as the average. Alternative promoters were tested (Supporting Information), but none showed both high stringency and inducibility and did not outperform OSQ7.3 for 2,3-oxidosqualene accumulation.

### Genome analysis of the 2,3-oxidosqualene overproducing strain

The degron strategy identified clone OSQ7.3 as capable of accumulating twice as much 2,3-oxidosqualen e as other clones of the same transformation. To determine possible genetic changes in this clone, genomic DNA from OSQ7.3 and OSQ7.2, a standard representative, was extracted and sequenced using next-generation sequencing. The individual short DNA segments were assembled with reference to the genome sequence of *S. cerevisiae* CEN.PK (NCBI sequence) in the CLC Genomics Workbench. Then, mutations of both strains were determined individually against this reference. Mutations common to OSQ7.2 and OSQ7.3 were masked to reveal those. These were nine insertions, one substitution, and one deletion (Fig. 2a). The only mutation in a coding sequence was found in the *ERG7* gene. Here, a single nucleotide deletion resulted in a frameshift and the formation of a stop codon 92 nucleotides downstream of the mutation and a chimera protein, extended by 28 amino acids compared to the native Erg7. The degron and associated sequences initially fused to *ERG7* were disrupted and not functional in this strain (Fig. 2b). This mutation was the only one detected with a 100% frequency in all short DNA segments across this sequence; all other mutations had a maximum frequency of 55%.

**Figure 2.**
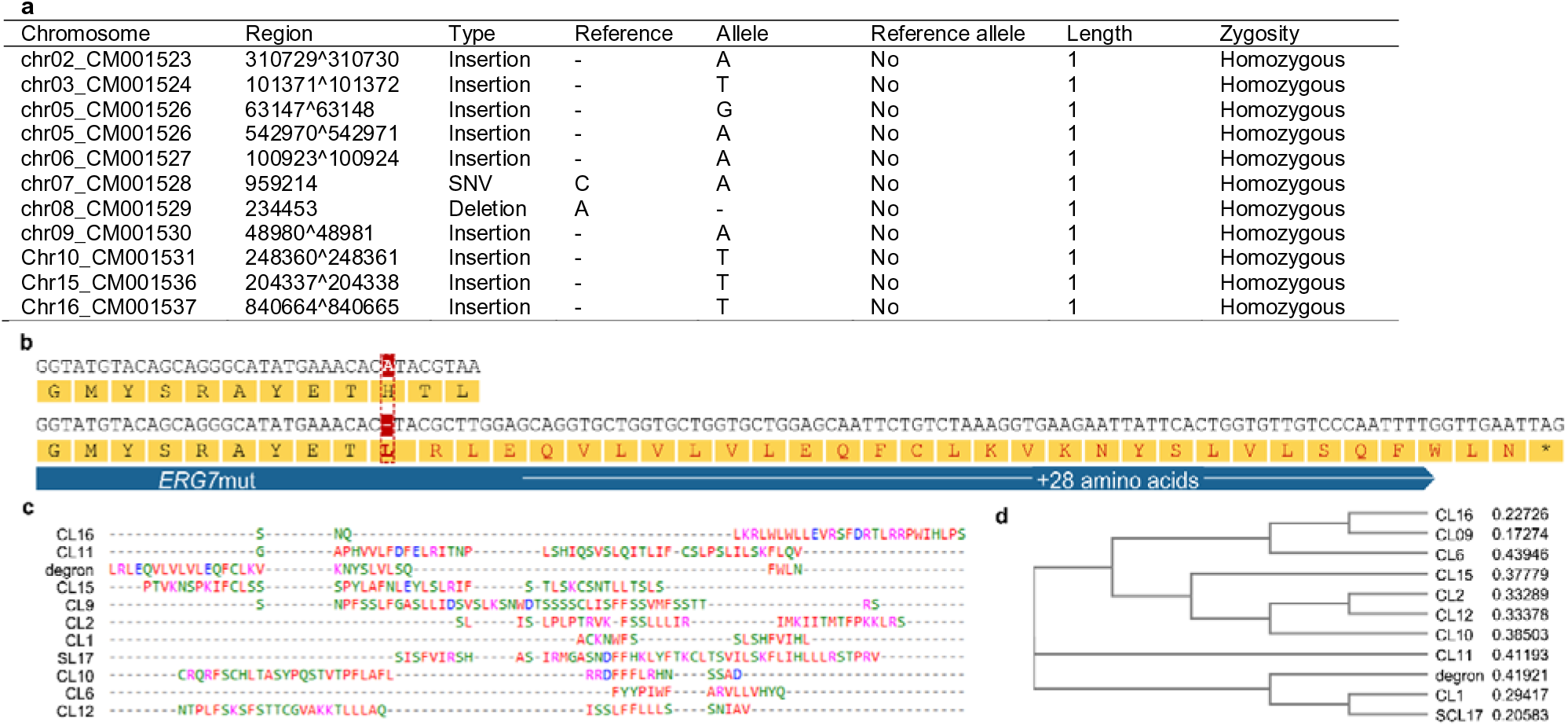
**(a)** Mutations specific for the 2,-oxidosqualene overaccumulatins clone OSQ7.3, **(b)** alignment of nucleotide and protein sequences of *ERG7* from CEN.PK and OSQ7.3, the single nucleotide deletion in the OSQ7.3 gene is highlighted in red **(c)** Alignment (CLUSTAL O (1.2.4) and **(d)** Neighbour joining tree of the *ERG7* degron tag and known degradation signals for ubiquitin system proteolysis.^1^

Multiple sequence alignment with known protein degradation signals revealed that the new degron tag is not similar to sequences known to destabilize proteins (Fig. 2c-d).^1^

### Reversed engineered Erg7p modification allows high OSQ accumulation

To validate that the mutated *ERG7* (*ERG7*mut) is responsible for the observed phenotype, we replaced the native gene in the reference strain CEN.PK 102-5B and further expressed *tHMG1* for deregulated MVA pathway activity (oSQ). We here refrained from deleting *ARE2* as in strain SQ1 to retrieve a lean engineering strategy and used an alternative POR, MTR1, which was shown to be superior for pentacyclic triterpenoid biosynthesis.^19^ The *ERG7* modification was introduced in strain rSQ by replacing 11 bp of the C-terminus (from the frameshift mutation to stop codon inclusive) with a cassette including the extended nucleotide sequence of the mutated *ERG7* (94 bp) followed by a hygromycin resistance cassette also present in OSQ7. The resulting strain, rOSQ, accumulated 2,3-oxidosqualene to 227 mg/L while the reference rSQ, expressing only *tHMG1*, solely accumulated squalene (137 mg/L squalene; 39 mg/g_DCW_) as observed before. We argued that the Erg7 modification results in either a reduced catalytic efficiency or reduced enzyme abundance due to decreased stability. To elucidate this further, we first quantified the sterol content in the yeast cells since reduced levels would support a reduced lanosterol synthase activity. Indeed, zymosterol and ergosterol content of rOSQ was significantly reduced compared to rSQ, strengthening our hypothesis of reduced activity. Contradicting, the lanosterol level was slightly increased in rOSQ, but measurement inaccuracy and underestimation of the measurement error might explain this apparent discrepancy. In addition, we run targeted proteomics on whole-cell extracts. This analysis could answer whether the proposed reduced flux into sterol biosynthesis is caused by reduced enzyme abundance and, therefore, activity on cell level. Whereas Erg7 was detected in the reference strain CEN.PK 102-5B, we failed to unambiguously identify Erg7mut in the engineered strain, indicating massively reduced abundance (data not shown).

Interestingly, strain rOSQ showed a significant growth defect and, as a result, a significantly increased specific titer of 2,3-oxidosqualene of 66 mg/g_DCW_, about 2-fold compared with strain OSQ7.3 (Fig. 1). A genetic difference between the two strains is the deletion of *ARE2* in OSQ7.3, while this sterol acyltransferase is present in rSQ and rOSQ. It is of now unknown how and if this contributes to the observed phenotypic differences. Another difference with OSQ7.3 is that the latter accumulates vast amounts of squalene in addition to 2,3-oxidosqualene, which together outpace the accumulation of 2,3-oxidosqualene in rOSQ.

### A lean engineering approach is sufficient for high triterpenoid production

With the 2,3-oxidosqualene overproducing strain in hand, we explored the potential to redirect this metabolite towards triterpenoid biosynthesis. As proof of concept, we chose the triterpenoid betulinic acid, derived from 2,3-oxidosqualene through cyclization to lupeol catalyzed by an oxidosqualene cyclase, which is then oxidized to betulinic acid through a single CYP/CPR couple via the corresponding alcohol and aldehyde (Fig. 3a). The enzyme screening ran parallel with the chassis development and used *S. cerevisiae* CEN.PK expressing only tHMG1 as a base. We initially evaluated diverse cyclases and revealed Olea europaea oxidosqualene synthase (OEW) as a promising monofunctional lupeol synthase (Fig. 3b). To identify CYPs capable of oxidizing lupeol into betulinic acid, we then coexpressed *OEW* and a cytochrome P450 reductase (POR) from *Lotus japonicus (LjCPR2)* with e*i*ght CYPs of subfamily CYP716A known to modify the C-28 position of triterpenoid scaffolds.^20^ However, only three enzymes produced low amounts of betulinic acid and larger amounts of alcohol betulin. Maximal conversio n of lupeol to alcohol, aldehyde, and acid was achieved with CYP716A15 (Fig. 3c).

**Figure 3.**
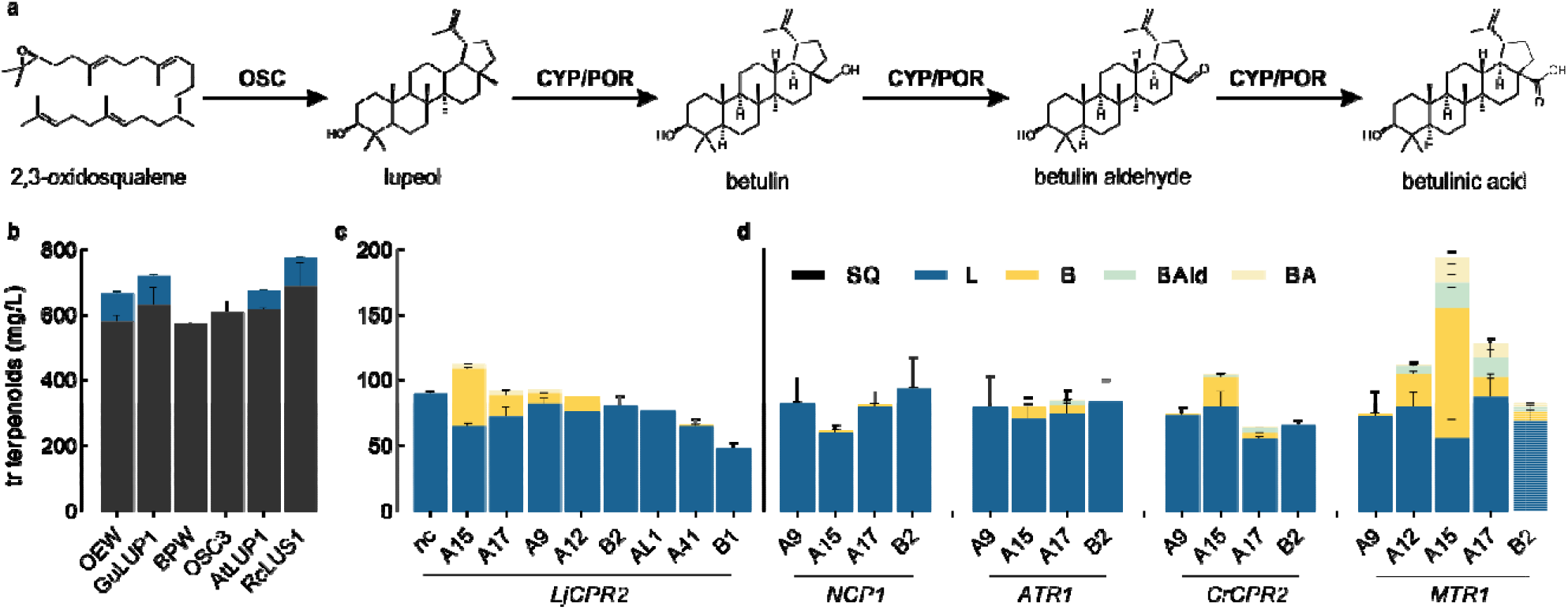
Combinatorial biosynthesis of betulinic acid biosynthesis (a): screening of oxidosqualene cyclases (b), cytochrome P450 monooxygenases (CYP) of family CYP716 (b), and cytochrome P450 reductases (POR) (c); SQ, squalene; L, lupeol; B, betulin; Bald, betulinic acid; BA, betulinic acid; nc, negative control (basis strain harboring an empty plasmid), (b-d), for enyzme abbreviations given on the x-axis see the supporting information.

CPRs mediate the essential electron transfer for Eukaryotic P450 activity. The promiscuous nature of the CPR electron donations allows for supporting several different P450s. However, P450/CPR pairing can significantly affect P450 catalytic activity and, in some cases, even alter P450 substrate promiscuity. General advice is to use the native CPR for optimal P450 interaction. However, this needs further consideration when working with P450s from plants, which can have multiple CPR isoforms with different activities or for which the native CP R is unknown.^21, 22^ We, therefore, tested different combinations of P450s and PORs, among which we identified CYP716A15/MTR as the best enzyme pair (Fig. 3d) Initial strain engineering efforts used the gene combination *LjCPR, CYP716A15*, and *OEW;* all genes were integrated into the genome of OSQ1.3 to create the betulinic acid-producing strain BA. For enhanced conversion, multiple copies of the cyclase and P450 genes were genomically integrated into retrotransposon Ty sites. In contrast, the CPR was integrated as a single copy to resemble the stoichiometric inferiority observed in natural systems because of the multiplexed interaction of the CPR with multiple P450s^23^ and because excess expression of POR leads to electron transfer uncoupling and the generation of toxic reactive oxygen species.^15^ BA6 produced 157.7 mg/L betulinic acid and a total triterpenoid titer of 1,345 mg/L in 98 hours of sake-flask cultivation in mineral salt medium with 50 g/L glucose, approximately 10-fold compared to the titers achieved in the nonoptimized strain during the initial screen (Table 1). The significant titers showcase the excellent potential of the chassis for triterpenoid biosynthesis. While betulin was again the main product of the P450 activity, lupeol was more efficiently converted into the oxidized triterpenoids, presumably because of the multi-copy integration of the P450 gene. The strain’s catalytic capabilities were further assessed in fed-batch fermentations using ethanol as the carbon source, previously identified to support high-titer betulinic acid production in yeast (Figure 4, Table 2).^24^

**Table 1.** Triterpenoid production in *S. cerevisiae* BA in batch shake-flask cultivation

| Time<br>h | CDW<br>g/L | Squalene<br>mg/L | Lupeol<br>mg/L | Betulin<br>mg/L | Betulin<br>aldehyde<br>mg/L | Betulinic acid<br>mg/L | Total triterpenoids<br>mg/L |
| --- | --- | --- | --- | --- | --- | --- | --- |
| 42 | 5.0 ± 0.3 | 40.2 ± 28 | 74.4 ± 3.2 | 157.2 ± 3.6 | 47.6 ± 0.5 | 86.2 ± 0.4 | 405.6 ± 7.1 |
| 98 | 11.1 ± 0.5 | 82.5 ± 1.8 | 301.3 ± 1.2 | 684.6 ± 18.4 | 119.4 ± 4 | 157.7 ± 2.1 | 1345.5 ± 5.5 |

**Figure 4.**
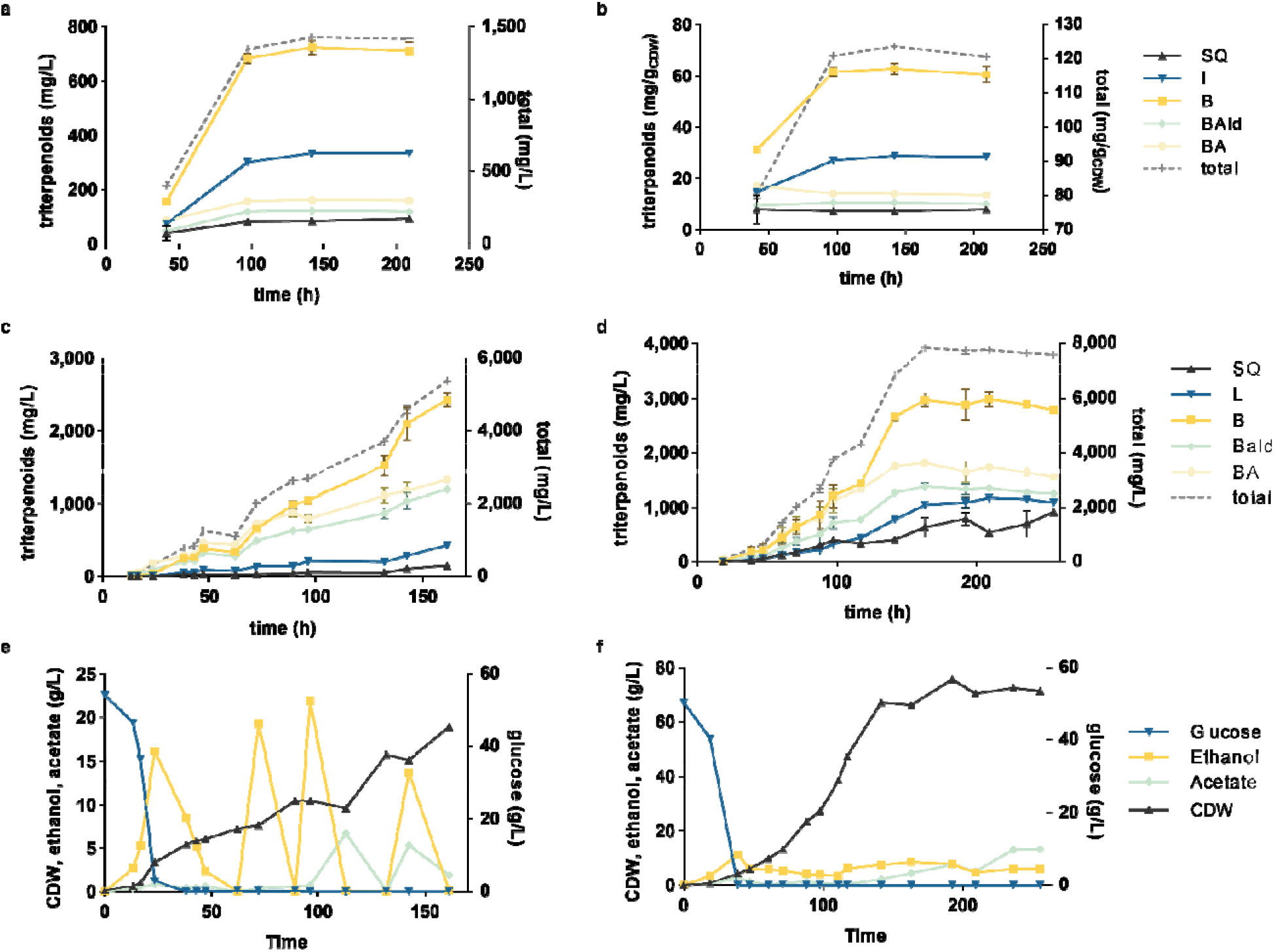
Product concentrations during fed-batch cultivation of *S. cerevisiae* BA in WM8+ medium. (a) Betulinic acid biosynthesis pathway. (b-c) triterpenoid titer in shake-flask cultivations (d,-g) Fermentation profiles of a DO-triggered ethanol pulse fed-batch; SQ-squalene, L-lupeol, B-betulin, BAld-betulinic aldehyde, BA-betulinic acid, EtOH-ethanol, Ac-acetate, CDW-cell dry weight, total triterpenoids corresponds to the sum of lupeol, betulin, betulin aldehyde, and betulinic acid.

**Table 2.** Volumetric concentrations and productivities of betulinic acid production in *S. cerevisiae* BA fed-batch cultivations with ethanol pulse feeds or controlled constant ethanol feed. Carbon content Cmol per pulse glucose or ethanol was equal. Total triterpenoid concentrations correspond to the sum of lupeol, betulin, betulin aldehyde, and betulinic acid.

| Mode | Time<br>h | Betulin |  | Betulinic acid |  | Total triterpenoids |  |
| --- | --- | --- | --- | --- | --- | --- | --- |
|  |  | mg/L | mgB/L/h | mg/L | mgBA/L/h | mg/L | mgTerp/L/h |
| EtOH pulse<br>(25 g/L) | 161 | 2432 | 15.1 | 1338 | 8.3 | 5400 | 33.5 |
| EtOH feed | 163 | 2967 | 18.2 | 1811 | 11.1 | 7194 | 44.1 |

Multi-copy gene integration into the Ty retrotransposon sites^25^ resulted in high lupane-type triterpenoid titers but turned out to be genetically unstable, especially under nonselective conditions in complex media (data not shown). Since robust and stable production is vital for an industrial process, we pursued an alternative integration method, in which the single integration sites are interspaced with essential genes so that destruction of expression cassettes through homologous recombination results in a lethal phenotype.^26^ Cyclase and P450 were the same as in OSQ1.3, but the POR from *Medicago truncatula* (*MTR1*) was used instead of the *L. japonicus* enzyme since *MTR1* had shown superior efficiency in lupeol oxyfunctionalization. Again the POR was integrated as a single copy, *OSC* and CYP were tested in single and double integrations (Fig. 5).

**Figure 5.**
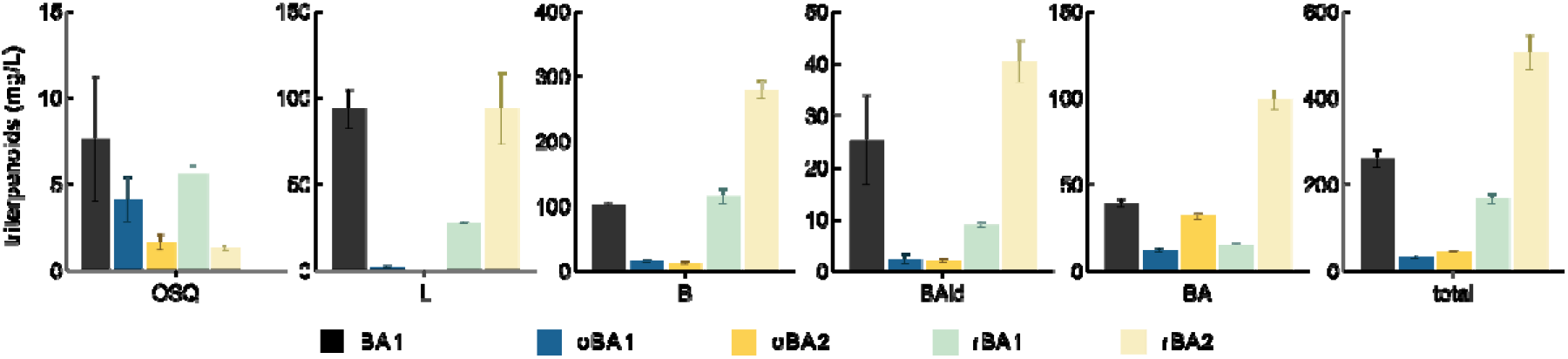
Production of lupane-type triterpenoids in strains expressing oxidosqualene cyclase OEW and CYP716A15 from single or double integrated gene copies (indicated by ‘1’ and ‘2’ in the strain name) in a background expressing the native *ERG7* (oBA1/2) or *ERG7*mut (BA1, oBA1/2. All strains were cultured in YPD medium with 2% glucose for 72h; error bars represent the standard deviation of three biological replicates.

**Figure 6.**
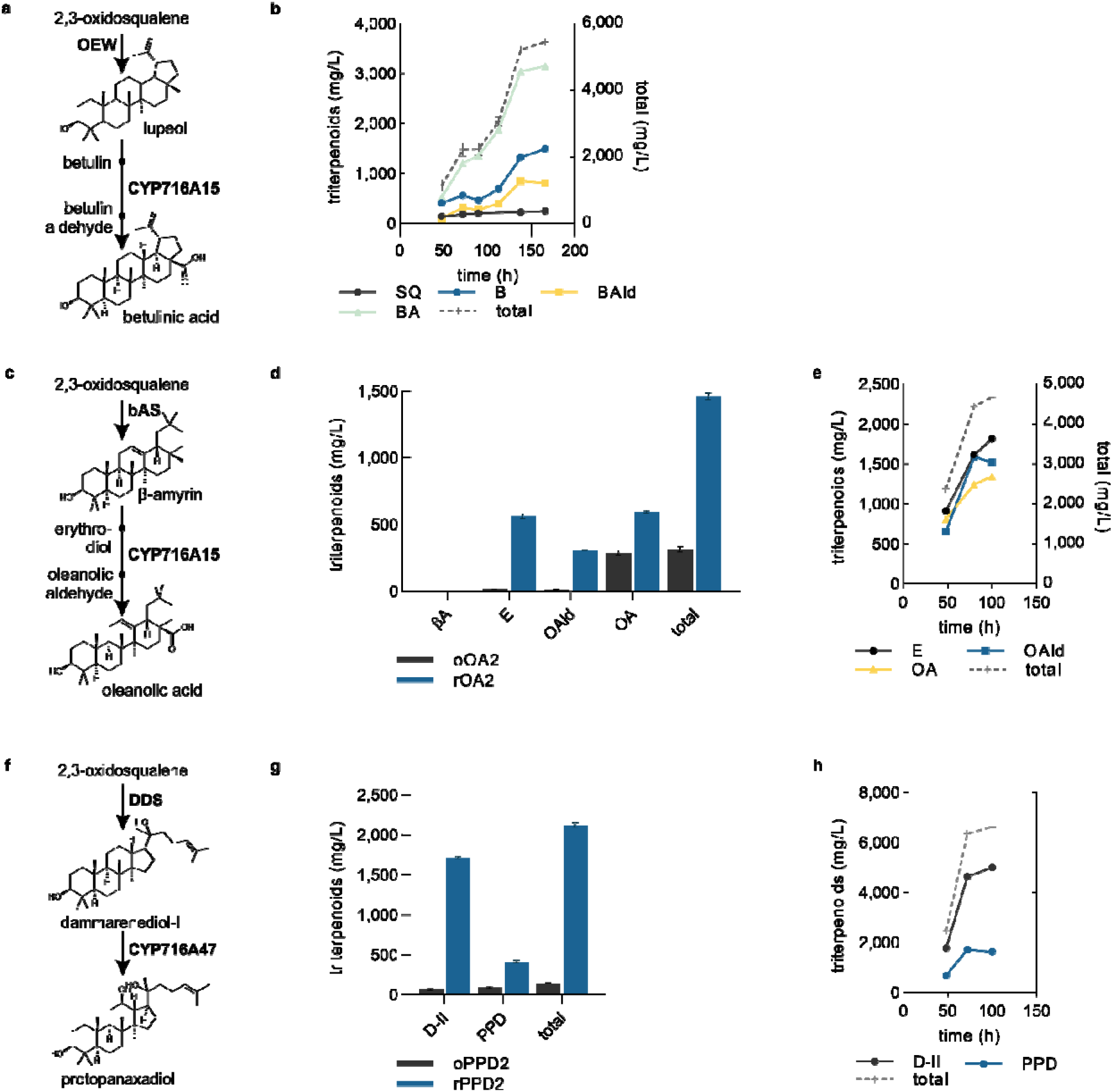
Capacity testing of rOSQ for the prodcution of betulinic acid (a), oleanolic acid (c) and protopanaxadiol (f). Comparison of triterpenoid accumulation in the rOSQ and rSQ background in shake-flask experiments (d,g; shake-flask data for betulinic acid production shown in Fig. 5); and performance in ethnaol fed-batch fermentation of rBA2 (b), rOA2 (e), and rPPD2 (h). SQ, squalene; B, betulin; Bald, betulin aldehyde; BA, betulinic acid; βA, β-amyrin; E, eryhtrodiol; Oald, oleanolic aldehyde; OA, oleanolic acid; D-II, dammarenediol-II; PPD, protopanaxadiol; total, sum of oxyfunctionalized triterpenoids. Error bars represent the standard deviation of biological replicates.

The triterpenoid production capabilities of the 2,3-oxidosqualene chassis were further tested on the pentacyclic oleanane-type triterpenoid oleanolic acid, the most comman agycone structures of saponins (glycosyalted triterepenoids) and the tetracyclic, dammarene-type triterpenoids, the ginsenoside precursors dammarenediol and protopanaxadiol. The same genetic engineering strategy were used, that is, single genomic integration of *MTR1* and double integration of the oxidosqualene cyclase and P450 encoding genes.

## Discussion

We present a general chassis for 2,3-oxidosqualene-dependent triterpenoid synthesis that supports high flux through the mevalonate pathway without heavy genome engineering. Using OSQ1.3 as chassis to produce triterpenoids enables the strain engineer to focus on the triterpenoid pathway, as abundant precursor supply is already provided. The novel mutation of Erg7 results in reduced lanosterol synthase abundance, allowing triterpenoid synthesis without abolishing sterol synthesis from the joint precursor 2,3-oxidosqualene.

While several studies aimed to reduce the flux towards sterols to facilitate flux rerouting into recombinant triterpenoid production pathways, the improvements in absolute terms from manipulating Erg7 activity were minor. Strategies ranged from full deletion to defined control of the transcription rate using repressible promoters for *ERG7* transcription or CRISPR interference.^13, 27-32^ Again, while the relative improvements compared to the reference were always impressive, the absolute amount of triterpenoids produced was most often modest.

The idea to use a degron to lower the half-life of Erg7p and thereby its abundance and activity was published this year, improving the production of the triterpenoid β-amyrin by 30% against the reference, and after further strain and process optimization contributed to 2.6 g/L β-amyrin production.^33^ The authors used a previously reported N-degron-mediated protein degradation mechanism ^34^ that was for example successfully installed for improving monoterpene synthesis in baker’s yeast.^35^. Briefly, the degron is first hydrolysed by the ubiquitin C-terminal hydrolase, exposing the N-end rule residue, ^36^ which subsequent leads to ubiquitination on the lysine-asparagine spacer, followed by protein degradation through the proteasome. The here presented degron relies also on the yeast protein quality assurance mechanisms and is independent of recombinant proteases. While the detailed mechanism was not investigated, the degron a. does not require interference by the experimenter, b. does not require an inductor, and c. does not distinguish between growth and production phase during the yeast culture, making its use very simple.

Comparing approaches to lower Erg7p activity is difficult, as no study exists where the different strategies were implemented in an identical strain background. We argue here that any undertaking lowering the competing flux to the sterols is ideally a. inducer-free, b., and allows good growth of the host, and c., most importantly, supports triterpenoid production. An explanation for the modest triterpenoid production in ergosterol auxotrophic yeast might be the requirement to supplement ergosterol into the medium for growth, which might trigger a low flux through the mevalonate pathway as the sterol is plentifully available. In the presence of ethanol, baker’s yeast incorporates more ergosterol into the cytoplasmic membrane, which is also a prerequisite for ethanol tolerance of yeast (see a recent review on the effect of low ergosterol levels on yeast phenotypes).^37^ While we used ethanol concentrations up to 25 g/L in the pulsed feed experiments, this falls considerable short of the 100+ g/L ethanol a yeast experiences during bioethanol production in industry. The used ethanol concentrations might have been, however, sufficient to activate the mevalonate pathway. In addition, low Erg7 abundance as here engineered and hence low ergosterol concentrations during growth on ethanol signal a high demand, explaining the comparably high flux through the mevalonate pathway we observed.^37-39^.

While our minimal engineering strategy is unique today, high titers of recombinantly produced triterpenoids, especially for the ginsenoside protopanaxadiol, with up to 10 g/L, were reported.^31, 40^ The authors optimized acetyl-CoA synthesis, overexpressed the entire mevalonate pathway, tinkered with *ERG7* expression, used ethanol as carbon source, and had biomass concentrations as high as 150 g_CDW_/L. While highly exciting that these high titers can be achieved, this strain and process optimization is not only very labor-intensive and hence hard to adopt by other groups, but it is also difficult to translate as biomass concentrations in industrial baker’s yeast fermentation do not exceed 60 to 80 g_CDW_/L because of limitations in oxygen supply besides others.

In conclusion, the present study shows that the simple engineering of *ERG7* combined with the classic HMG1 truncation (tHmg1) is sufficient for abundant 2,3-oxidosqualene availability and allows effective rerouting of the carbon flux into recombinant triterp biosynthesis. The chassis presented makes it simple to explore the triterpenoids, this intriguing group of natural products for the greater good, e.g., improving food, cosmetic, and pharmaceutical applications, while reducing the pressure on the plants and other possibly endangered organisms from which triterpenoids originate.

## Methods

### Strain construction

Strains, plasmids, and primers used are listed in Supplementary Tables S1-S3. If not stated otherwise, *S. cerevisiae* CEN.PK 102-5B was used as the parent strain. All heterologous genes were codon-optimized for *S. cerevisiae* and ordered from GeneArt (Thermo Fisher Scientific Inc., USA), GenScript (New Jersey, USA), and Integrated DNA Technology (Leuven, Belgium) as given in Supplementary Table S5.

All plasmids were constructed based on EasyClone vectors.^25, 26^ Digested vectors were gel purified and integrative vectors assembled with a classical uracil excision cloning reaction.^41^ The genes encoding for the P450 reductase from *Medicago truncatula* (MTR1), dammarenediol synthase (PgDDS), and protopanaxadiol synthase (CYP716A47) from *Panax ginseng* were synthesized by Integrated DNA Technologies (Leuven, Belgium) as codon-optimized gBlocks and PCR amplified with primer pairs p7, p8, and p10, respectively. Fragment *P*_*PGK1*_*-P*_*TEF1*_*-tHMG1* was amplified from pCfB_0825 using primer pair 1. Plasmid pESC-AaBAS, carrying the β-amyrin synthase from *Artemisia annua*, was kindly provided by Prof. J. D. Keasling (UC Berkeley, Berkeley, CA, USA).^29^ The AaBAS gene was amplified from pESC-AaBAS using primer pair p6. Correct amplification of DNA fragments was confirmed by diagnostic PCR and Sanger sequencing. The EasyClone vectors pCfB255, pCfB257, pCfB258, pCfB259, and pCfB390 were digested with AsiSI and subsequently nicked with Nb.BsmI. Fragments P_*TEF1*_–*tHMG1* and *P*_*PGK1*_*-*MTR were cloned into pCfB259, resulting in p-2647-tm. Fragment OEW–*P*_*PGK1*_–P_*TEF1*_–CYP716A15, P_PGK1_-AaBAS-P_TEF1_-CYP716A15, and P_PGK1_-PgDDS-P_TEF1_-CYP716A47 were respectively ligated into pCfB390 and pCfB258, resulting in plasmids p-2652-LP, p-2646-LP, p-2652-BP and p-2646-BP, p-2652-PP and p-2646-PP.

Integrative vectors were digested by NotI and gel purified. Using the lithium acetate transformation protocol, *S. cerevisiae* strains were transformed with circular or linearized plasmids.^42^ transformants were selected on YPD plates supplemented with 200 mg/L hygromycin B or drop-out agar media. Approximately 1 μg DNA of each vector was transformed into competent yeast cells, and transformants were selected on drop-out agar media.

Strain TC0 was constructed by integrating t*HMG1* and MTR1 genes into *S. cerevisiae* CEN.PK 102-5B. Strain TC1 was constructed by mutating the *ERG7* gene in strain TC0. To mutate the *ERG7* gene, the hphMX cassette was amplified from plasmid pSH69 using primer pair p10 and transformed into strain TC0. Strains TC0-LP and TC1-LP were constructed by integrating OEW and CYP716A15 genes into TC0 and TC1. Strains TC0-2LP and TC1-2LP were constructed by integrating OEW and CYP716A15 genes into TC0-LP and TC1-LP. TC0-2BP and TC1-2BP were constructed by sequentially integrating two copies of AaBAS and CYP716A15 genes into TC0 and TC1. TC0-2PP and TC1-2PP were constructed by sequentially integrating two copies of PgDDS and CYP716A47 genes into TC0 and TC1. All transformants were verified by diagnostic PCR.

### Cultivation conditions

*Escherichia coli* was propagated in Lysogeny Broth (LB) medium, complemented with 100 mg/mL ampicillin for selection and plasmid maintenance where appropriate. *S. cerevisiae* strains were cultivated in YPD medium (per liter: glucose 20 g, yeast extract 10 g, peptone 20 g) or WM8+ mineral salt medium (per liter: (NH_4_)H_2_PO_4_ 0.25 g, NH_4_Cl 2.80 g, sodium glutamate 10 g, MgCl_2_·6H_2_O 0.25 g, CaCl_2_·2H_2_O 4.60 g, KH_2_PO_4_ 2 g, MgSO_4_·7H_2_O 0.55 g, myoinositol 100 mg, ZnSO_4_·7H_2_O 6.25 mg, FeSO_4_·7H_2_O 3.50 mg, CuSO_4_·5H_2_O 0.40 mg, MnCl_2_·4H_2_O 0.10 mg, MnCl_2_·2H_2_O 1.00 mg, Na_2_MoO_4_·2H_2_O 0.50 mg, nicotinic acid 11.00 mg, pyridoxin-HCl 26.00 mg, thiamin-HCl 11.00 mg, biotin 2.55 mg, calcium pantothenate 51 mg, Na_2_EDTA 2.92 g, CoCl_2_.6H_2_O 0.30 mg, H_3_BO_3_ 1.00 mg, KI 0.10 mg, and *p⍰*aminobenzoic acid 0.20 mg.^24^ *S. cerevisiae* strains with multi-copy integrations of heterologous genes into Ty retrotransposon sites were strictly maintained on/in mineral salt media to avoid loss of gene copies under nonselective conditions.^25^ All shake flask experiments (500 mL flasks containing 50 mL medium (10 % (v/v))) in rotary shakers at 30 °C, 300 rpm and 5 cm amplitude. A 24-h pre-culture grown under the same conditions was used to inoculate the main cultures to a start OD_600_ of 0.1 or 0.4.

Samples were taken periodically to monitor growth and product formation. Optical densities at 600 nm (OD_600_) were measured spectrophotometrically using a Ultrospec 10 cell density meter (Amersham Biosciences, Glattbrugg, Schweiz). Cell dry weights (CDW) were either calculated by multiplying the OD_600_ signal with an experimentally pre-determined correlation factor of 0.21 g/L per unit OD_600_ or gravimetrically, as reported previously.^24^

### Triterpenoid and sterol quantification

Triterpenoids were quantified by HPLC-CAD following a published protocol.^43^ Specifically, samples of 800 μL of culture broth were transferred into 2 mL sample tubes to analyze intracellular triterpenoid concentrations and were stored at −20 °C until further processing or immediately analyzed. Cells were disrupted by adding 250 μL of glass beads (smooth 500 μm, Carl Roth, Karlsruhe, Germany), 80 μL 1 M HCl, and 800 μL chloroform: methanol (ratio 80:20, v/v) and vigorously agitating in the Mini-Beadbeater 16 (BioSpec Products, Bartlesville, USA) for 4 min. Samples were centrifuged for 10 min at 16,089 x g and 5 °C. 200 μL of the lower organic phase were transferred into a glass vial (VVial, clear, 9 mm, screw, Phenomenex) containing an insert (15 mm tip, wide opening, 0.2 mL, 6×31 mm, clear, Macherey-Nagel GmbH & Co. KG Germany) for High-Performance Liquid Chromatography (HPLC) analysis. Product quantification was conducted with a reverse HPLC system (Thermo Fisher UltiMate 3000 Dionex). Samples were kept a 4 °C in an autosampler (Thermo Fisher UltiMate 3000). Five μL of the sample were injected into a C18 column (Merck Spherisorb ODS-2 (5 μm), 250×4 mm, Merck KGaA, Germany) with a guard column (EC 4/3 NUCLEODUR C18 Gravity, 3 μm, Macherey-Nagel GmbH & Co. KG Germany), while the column temperature was kept at 40 °C in a thermostatic column compartment (Thermo Fisher UltiMate 3000). Analytes were eluted using gradient chromatography at 1.2 mL/min flow rate using a binary pump (Ultimate 3000 Pump, Dionex). Solvent A was acetonitrile (LC-MS grade). Solvent B was 0.2 % (v/v) formic acid of high purity (≥ 98 %). The linear gradient method was as follows: 20 % B from 0-1 min, 20-0 % B from 1-13 min, 0 % B held until 23 min, then back to 20 % B from 23-26 min. Solvents were degassed using an online degasser. Analytes of interest were monitored using a charged aerosol detector (CAD, esa ERC GmbH) driven by a nitrogen generation system (Parker, BALSTON, Analytical Gas Systems). Technical duplicates were measured, i. e., two times 800 μL were taken from the same cultivation, processed and analyzed. For quantification of triterpenoids, serial dilutions (df = 2) of standards for squalene (Mo Bi Tec, Göttingen Germany, purity >95%), 2,3-oxidosqualene (both from MoBiTec (Göttingen Germany, ≥97.5%), lupeol (Sigma Aldrich, ≥ 94%), betulin (Sigma Aldrich, ≥ 98%), betulinic acid (Sigma Aldrich, ≥ 98%), betulin aldehyde (Cfm Oskar Tropitzsch GmbH, Marktredwitz Germany, ≥ 98%), β-Amyrin (Biomol, Hamburg Germany, >95%), erythrodiol (Sigma Aldrich, ≥97%), oleanolic acid (Sigma Aldrich, ≥98%), and protopanaxadiol (Sigma Aldrich, ≥96%) in chloroform were measured.

Due to the lack of authentic standards of oleanolic aldehyde and dammarenediol-II, oleanolic aldehyde and dammarenediol-II were first identified by gas chromatography-mass spectrometry (C_MS) and Liquid chromatography-MS and then quantified by HPLC-CAD as described above using the response curves of betulinic aldehyde and protopanaxadiol, respectively, as proxies.^44^

GC-MS analysis was carried out on a Trace GC Ultra gas chromatograph equipped with a TSQ single quadrupole mass spectrometer (Thermo Scientific, Waltham, MA, USA). The temperature of the ion source equalled to 290 °C. Electron impact (EI) ionization was kept at 70 eV, and a constant flow rate of 1 mL/min helium was applied. The injection volume for every sample was 1 μL and the injector temperature was set to 250 °C. The temperature was initially held at 80 °C for 1 min and was increased with a gradient of 20 °C/min to 320 °C, held for 6 min.

LC-MS analyses were carried out on an Agilent 1200 system (Agilent Technologies, Santa Clara, USA) coupled to an LTQ Orbitrap XL (Thermo Fisher Scientific Inc., Waltham, USA) equipped with an atmospheric pressure chemical ionization (APCI) source used in positive ion mode under the following conditions: source voltage 3 kV, source current 5.8 μA, vaporizer temperature 400°C, capillary voltage 4 V, capillary temperature 275 °C. Compounds were separated on a C18 column (Merck Spherisorb ODS-2 (5 μm), 250×4 mm, Merck KGaA, Germany) using an acetonitrile-water (containing 0.2 % (v/v) formic acid) gradient at a flow rate of 1 mL/min. Chromatographic separation remained the same as used for HPLC-CAD analysis.

For lanosterol, zymosterol and ergosterol quantification, the samples were prepared based on a previously described procedure with some modifications.^45, 46^ 2 mL cell suspensions were spun down; the supernatant was discarded, and the pellet was resuspended in 0.4 mL KOH (20% (wt/vol) in 50% ethanol) containing 10 μg/mL cholesterol as internal standard and transferred into a 2 mL screw-cap tube. To ensure complete saponification, the sample tube was placed into boiling water for 5 min. Glass beads (0.5mm) and 0.4 mL of hexane were added, and cells were disrupted by vortexing vigorously for 5 minutes. The tubes were centrifuged for 10 min at 13,000 rpm, and the top hexane layer was transferred to a 1.5 mL GC-vial and evaporated in a centrifugal evaporator (SpeedVac SC100, Savant, New York, NY) for 20 min at 50 °C. Finally, dried sterols were resuspended in 200 μL chloroform. For the GC-FID analysis, 30 μL of the organic phase was transferred in a 1.5 mL GC-vial mixed with 30 μL MSTFA (N-Methyl-N-(trimethylsilyl) trifluoroacetamide), vortexed again and derivatized for 30 min at 70 °C in the heating block.

Sterol quantification was achieved with a Trace GC Ultra (Thermo Scientific, USA) equipped with a flame ionization detector (GC-FID) and a Trace GOLD TG-5SilMS capillary column (length, 15 m; inner diameter, 0.25 mm; film thickness, 0.25 μm) at a constant nitrogen flow of 1 mL/min. The injection volume for every sample was 1 μL. The GC oven temperature was initially set to 100 °C for 1 min, followed by an increase to 320 °C at a rate of 20 °C/min, which was finally maintained for 6 min. For quantification, serial dilutions (df = 2) of standards for lanosterol (Biomol (Hamburg Germany, purity >95%), zymosterol (Sigma Aldrich, Taufkirchen, Germany, purity >99%), and ergosterol (Sigmal Aldrich, Taufkirchen, Germany, purity ≥97.5%) in chlorofom were measured.

To quantify carbon sources and extracellular products (glucose, ethanol, glycerol and acetate), samples were taken at regular time points. Biomass was harvested by centrifugation; the supernatant was filtered (0.22 μm, syringe filter) and transferred into HPLC vials (VVial, clear, 9 mm, screw, Phenomenex). Extracellular metabolites were measured using high-performance liquid chromatography (HPLC) with UV and RI detection. A polystyrol-divinylbenzene copolymer column (PS-DVB, CS-Chromatographie, Germany) was used, and analytes were eluted with freshly prepared running buffer (5 mM H_2_SO_4_) at 0.6 mL/min at 30 °C for 30 min. The sample injection volume amounted to 20 μL. For quantification of metabolites, serial dilutions (df = 2) of standards for D-(+)-glucose, ethanol, glycerol, and acetate (all from Sigma-Aldrich (Taufkirchen, Germany), at a purity of ≥99%) were prepared in a range between 0.5 g/L to 50 g/L.

## Acknowledgments

We acknowledge financial support by the Deutsche Bundesstiftung Umwelt (Grant ID AZ13272). HG acknowledges the financial support by a personal stipend of the Chinese Scholarship Council (CSC). pSM1959 was a gift from Susan Michaelis (Addgene plasmid # 41837; http://n2t.net/addgene:41837; RRID: Addgene_41837).

## Supporting Information

**Table S3**. Strains used in this study

**Table S4**. Plasmids used in this study

**Table S5**. Primers used in this study

**Table S6**. Triterpenoid production in engineered yeast (combinatorial biosynthesis)

**Table S7**. Synthetic genes used in this study

