## Supporting Information for "Triterpenoid production with a minimally engineered *Saccharomyces cerevisiae* chassis"

Table S1. Strains used in this study

| **Strain ID** | **Genotype; relevant genetic modifications** | **Description** | **Reference/ Origin** |
| --- | --- | --- | --- |
| DH10B | *Δ(ara-leu) 7697 araD139 fhuA ΔlacX74 galK16 galE15 e14-ϕ80dlacZΔM15 recA1 relA1 endA1 nupG rpsL (StrR) rph spoT1 Δ(mrr-hsdRMS-mcrBC)* | Cloning strain | New England Biolabs |
| CEN.PK2 | MATα*; ura3-52; leu2-3_112; TRP1; his3delta1; MAL2-8C; SUC2* | Parent strain for combinatorial testing of gene candidates | Euroscarf |
| CEN.PK | MATa*, ura3-52, his3-11, leu2-3/112, TRP1, MAL2-8c, SUC2* | Parent strain for metabolic engineering | Euroscarf |
| SQ1 | CEN.PK 102-5B, *tHMG1,* ATR2, *are2::*KanMX | Parent strain for testing Erg7 degron strategy | ^1^ |
| CK24 | SQ1 + P*_CUP1_*-3vGFP | Evaluation of inducible and conditionally active promoters to control TEV protease expression | This work |
| CK25 | SQ1 + P*_AHD2_*-3vGFP |  | This work |
| CK26 | SQ1 + P*_HTX7_*-3vGFP |  | This work |
| CK28 | SQ1 + P*_CUP1_-SPO13*-3vGFP |  | This work |
| OSQ6 | SQ1 + erg7::*ERG7*tag | Targeted degradation of Erg7p | This work |
| OSQ7 | OSQ6 + P*_CUP1_* controlled TEV protease |  | This work |
| OSQ7.1 | OSQ7 clone 1 |  | This work |
| OSQ7.2 | OSQ7 clone 2 |  | This work |
| OSQ7.3 | OSQ7, frameshift mutation in *ERG7* overrides stop codon |  | This work |
| OSQ8 | OSQ6 + P*_ADH2_* controlled TEV protease |  | This work |
| OSQ9 | OSQ6 + P*_HXT7_* controlled TEV protease |  | This work |
| OSQ10 | OSQ6 + P*_CUP1-SPO13_* controlled TEV protease |  | This work |
| OSQ11 | SQ1 + P*_ERG7_* replaced with p*_HXT1_* | Transcriptional control of *ERG7* expression | This work |
| rSQ | CEN.PK 102-5B, single-copy integration of T*_ADH1_–tHMG1–*P*_TEF1_–*P*_PGK1_*–*MTR–*T*_CYC1_* cassettes and *KlLEU2* marker gene into chromosome X |  | This work |
| rOSQ | rSQ with replacement of the native *ERG7* gene with the frame-shift *ERG7* mutant |  | This work |
| BA | Simo1575, multi-copy genomic integration of T*_CYC1_–OEW*–P*_PGK1_–*P*_TEF1_–CYP716A15*–T*_ADH1_* cassette and *KlURA3* marker gene into retrotransposon sites Ty4cons | Betulinic acid production | This work |
| oBA1 | rSQ, single-copy integration of T*_CYC1_–OEW–*P*_PGK1_*–*P_TEF1_*–CYP716A15*–*T*_ADH1_* cassettes and *URA3* marker gene integrated into chromosome XI |  | This work |
| rBA1 | rOSQ, single-copy integration of T*_CYC1_–OEW–*P*_PGK1_*–*P_TEF1_*–CYP716A15*–*T*_ADH1_* cassettes and *URA3* marker gene integrated into chromosome XI |  | This work |
| BA1 | Simo1575, single-copy genomic integration of T*_CYC1_–OEW*–P*_PGK1_–*P*_TEF1_–*CYP716A15–T*_ADH1_* cassette and *URA3* marker gene into chromosome XI |  |  |
| oBA2 | rSQ-LP, T*_CYC1_–OEW–*P*_PGK1_*–*P_TEF1_*–CYP716A15*–*T*_ADH1_* cassettes and *HIS3* marker gene integrated into chromosome XII of |  | This work |
| rBA2 | rOSQ-LP, T*_CYC1_–OEW–*P*_PGK1_*–*P_TEF1_*–CYP716A15*–*T*_ADH1_* cassettes and *HIS3* marker gene integrated into chromosome XII |  | This work |
| oOA2 | rSQ, T*_CYC1_*–AaBAS–P*_PGK1_*–P*_TEF1_*–CYP716A15–T*_ADH1_* cassettes and *URA3* marker gene, T*_CYC1_*–AaBAS–P*_PGK1_*–P*_TEF1_–*CYP716A15–T*_ADH1_* cassettes and *HIS3* marker gene sequentially integrated into chromosome XII |  | This work |
| rOA2 | rOSQ, T*_CYC1_*–AaBAS–P*_PGK1_*–P*_TEF1_–*CYP716A15–T*_ADH1_* cassettes and *URA3* marker gene, T*_CYC1_*–AaBAS–P*_PGK1_*–P*_TEF1_–*CYP716A15–T*_ADH1_* cassettes and *HIS3* marker gene sequentially integrated into chromosome XII |  | This work |
| oPPD2 | rSQ, T*_CYC1_*–PgDDS–P*_PGK1_*–P*_TEF1_*–CYP716A47–T*_ADH1_* cassettes and *URA3* marker gene, T*_CYC1_*–PgDDS–P*_PGK1_*–P*_TEF1_*–CYP716A47–T*_ADH1_* cassettes and *HIS3* marker gene sequentially integrated into chromosome XII |  | This work |
| rPPD2 | rOSQ, T*_CYC1_*–PgDDS–P*_PGK1_*–P*_TEF1_*–CYP716A47–T*_ADH1_* cassettes and *URA3* marker gene, T*_CYC1_*–PgDDS–P*_PGK1_*–P*_TEF1_*–CYP716A47–T*_ADH1_* cassettes and *HIS3* marker gene integrated into chromosome XII |  | This work |
| OBref | CEN.PK111-61A, *ura3::tHMG1, kanMX* | Upregulated MVA pathway | This work |
| OB1 | CEN.PK2U *OEW* (episomal expression from CEN/ARS plasmid) | Screening of oxidosqualene cyclases | This work |
| OB2 | CEN.PK2U *GuLUP1* (episomal expression from CEN/ARS plasmid) |  | This work |
| OB3 | CEN.PK2U *BPW* (episomal expression from CEN/ARS plasmid) |  |  |
| OB4 | CEN.PK2U *OSC3* (episomal expression from CEN/ARS plasmid) |  | This work |
| OB5 | CEN.PK2U *AtLUP1*(episomal expression from CEN/ARS plasmid) |  | This work |
| OB6 | CEN.PK2U *RcLUS1* (episomal expression from CEN/ARS plasmid) |  | This work |
| OB7 | CEN.PK2U, episomal expression of *OEW*, *LjCPR1*, *CYP716A12* from CEN/ARS plasmids | Screen of cytochrome P450 monooxygenases | This work |
| OB8 | CEN.PK2U, episomal expression of *OEW*, *LjCPR1*, *CYP716AL1* from CEN/ARS plasmids |  | This work |
| OB9 | CEN.PK2U, episomal expression of *OEW*, *LjCPR1*, *CYP716A15* (episomal expression from CEN/ARS plasmid) |  | This work |
| OB10 | CEN.PK2U, episomal expression of *OEW*, *LjCPR1*, *CYP716A17* (episomal expression from CEN/ARS plasmid) |  | This work |
| OB11 | CEN.PK2U, episomal expression of *OEW*, *LjCPR1*, *CYP716A9* (episomal expression from CEN/ARS plasmid) |  | This work |
| OB12 | CEN.PK2U, episomal expression of *OEW*, *LjCPR1*, *CYP716AB1* (episomal expression from CEN/ARS plasmid) |  | This work |
| OB13 | CEN.PK2U, episomal expression of *OEW*, *LjCPR1*, *CYP716A41* (episomal expression from CEN/ARS plasmid) |  | This work |
| OB14 | CEN.PK2U, episomal expression of *OEW*, *LjCPR1*, *CYP716AB2* from CEN/ARS plasmids |  | This work |
| OB15 | CEN.PK2U, episomal expression of *OEW*, *ATR1*, *CYP716A15* from CEN/ARS plasmids | Screen of cytochrome P450 reductases | This work |
| OB16 | CEN.PK2U, episomal expression of *OEW*, *ATR1*, *CYP716A17* from CEN/ARS plasmids |  | This work |
| OB1 | CEN.PK2U *OEW*, CEN.PK2U, episomal expression of *ATR1*, *CYP716A9* from CEN/ARS plasmids |  | This work |
| OB18 | CEN.PK2U episomal expression of *OEW*, *NCP1*, *CYP716A15* from CEN/ARS plasmids |  | This work |
| OB19 | CEN.PK2U episomal expression of *OEW*, *NCP1*, *CYP716A17* from CEN/ARS plasmids |  | This work |
| OB20 | CEN.PK2U episomal expression of *OEW*, *NCP1*, *CYP716A9* from CEN/ARS plasmids |  | This work |
| OB21 | CEN.PK2U episomal expresion of *OEW*, *CrCPR*, *CYP716A15* from CEN/ARS plasmids |  | This work |
| OB22 | CEN.PK2U CEN.PK2U, episomal expression of *OEW*, *CrCPR*, *CYP716A17* from CEN/ARS plasmids |  | This work |
| OB23 | CEN.PK2U, episomal expression of *OEW*, *MTR1*, *CYP716A15* from CEN/ARS plasmids |  | This work |
| OB24 | CEN.PK2U CEN.PK2U, episomal expression of *OEW*, *CrCPR*, *CYP716A9* from CEN/ARS plasmids |  | This work |
| OB25 | CEN.PK2U episomal expression of *OEW*, *MTR1*, *CYP716A17* from CEN/ARS plasmids |  | This work |
| OB26 | CEN.PK2U episomal expression of *OEW*, *LjCPR1*, *CYP716A9* from CEN/ARS plasmids |  | This work |
| OB27 | CEN.PK2U episomal epxression of *OEW*, *MTR1*, *CYP716A9* from CEN/ARS plasmids |  | This work |

Table S2. Plasmids used in this study

| **Plasmid** | **Relevant description** | **Reference** |
| --- | --- | --- |
| pSH69 | *P_AgTEF1_-hphMX-T_ScCYC1_*, clonNAT resistance, Amp^R^ | Euroscarf |
| pUG6 | Template for KanMX cassette used as marker in gene deletion | Euroscarf |
| pUC57 | Cloning vector | GenScript |
| pTT | *E. coli*-yeast shuttle vector (colE1 ori, AmpR; CEN/ARS) | Organobalance GmbH |
| pESC-AaBAS | Cloning *P_GAL10_-AaBAS-T_ADH1_* cassette into pESC | ^2^ |
|  | Integrative vector, genomic integration into chromosomes XII-1 sites, Amp^R^, USER cassette, *KlLEU2* | ^3^ |
| pCfB257 | Integrative vector, genomic integration into chromosomes X-3 sites, Amp^R^, USER cassette, *KlLEU2* | ^3^ |
| pCfB258 | Integrative vector, genomic integration into chromosomes X-4 sites, Amp^R^, USER cassette, *SpHIS5* | ^3^ |
| pCfB259 | Integrative vector, genomic integration into chromosomes XII-1 sites, Amp^R^, USER cassette, *KlLEU2* | pCfB259 |
| pCfB390 | Integrative vector, genomic integration into chromosomes XI-3 sites, Amp^R^, USER cassette, *KlURA3* | ^3^ |
| pCfB2796 | vector for multiple integrations at sites sharing homology with Ty4Cons, Amp^R^, USER cassette, *KlURA3* | ^4^ |
| pCfB_0825 | pCfB257 with T*_ADH1_–tHMG1–* P*_TEF1_-*P*_PGK1_–ATR2–*T*_CYC1_* insert | ^1^ |
| pCfB_0719 | pCfB322 with T*_ADH1_–CYP716AL1–* P*_TEF1_-*P*_PGK1_–AtLUP1–*T*_CYC1_* insert | ^1^ |
| pSM1959 | pRS425 with *SEC63*-mRFP insert, ori(pMB1), amp^R^ , 2µ, *LEU2* selection marker | ^5^, AddGene (plasmid #41837) |
| p2796_BP | Cloning of *P_PGK1_–OEW–T_CYC1_*, *P_TEF1_–CYP716A15–T_ADH1_* cassettes into pCfB2796 | This work |
| p2652-*OEW* | Cloning of *P_PGK1_–OEW–T_CYC1_* cassettes into pCfB390 | This work |
| p2647-tm | Cloning of *P_TEF1_–tHMG1–T_ADH1_*, *P_PGK1_–MTR1–T_CYC1_* cassettes into pCfB259 | This work |
| p2647-ta | Cloning of *P_TEF1_–tHMG1–T_ADH1_*, *P_PGK1_–ATR2–T_CYC1_* cassettes into pCfB259 | This work |
| p2652-LP | Cloning of *P_PGK1_–OEW–T_CYC1_*, *P_TEF1_–CYP716A15–T_ADH1_* cassettes into pCfB390 | This work |
| p2646-LP | Cloning of P_PGK1_–*OEW*–T_CYC1_, P_TEF1_–CYP716A15–T_ADH1_ cassettes into pCfB258 | This work |
| p2652-BP | Cloning of *P_PGK1_–AaBAS–T_CYC1_*, *P_TEF1_–CYP716A15–T_ADH1_* cassettes into pCfB390 | This work |
| p2646-BP | Cloning of *P_PGK1_–AaBAS–T_CYC1_*, *P_TEF1_–CYP716A15–T_ADH1_* cassettes into pCfB258 | This work |
| p2652-PP | Cloning of *P_PGK1_–PgDDS–T_CYC1_*, *P_TEF1_–CYP716A47–T_ADH1_* cassettes into pCfB390 | This work |
| p2646-PP | Cloning of *P_PGK1_–PgDDS–T_CYC1_*, *P_TEF1_–CYP716A47–T_ADH1_* cassettes into pCfB258 | This work |
| p2652-*OEW* | Cloning of *P_PGK1_–OEW–T_CYC1_* cassettes into pCfB390 | This work |

Table S3. Primers used in this study

| **Primer ID** | **Primer name** | **Sequence** | **Application/Ref.** |
| --- | --- | --- | --- |
| p1_fw | tHMG1_fw | CGTGCGAUTCAGGATTTAATGCAGGTGACGGA | *tHMG1* amplification |
| p1_rv | tHMG1_fw | AGTGCAGGUAAAACAATGACTGCAGACCAATTGGTG | tHMG1 amplification |
| p2_fw | AtATR2_fw | ATCTGTCAUAAAACAATGTCCTCCTCTTCTTCATCATCCACC | ^1^ |
| p2_rv | AtATR2_rv | CACGCGAUTCACCAGACATCTCTCAA | ^1^ |
| p3_fw | pTEF1_fw | ACCTGCACUTTGTAATTAAAACTTAGATTAGATTGCTATGC | P*_TEF1_* amplification |
| p3_rv | pPGK1_rev | ATGACAGAUTTGTTTTATATTTGTTG | ^1^ |
| p3_rv2 | pPGK1_fw | CACGCGAUGGAAGTACCTTCAAAGAATGGGGTC | P*_PGK1_*-*OEW*–T*_CYC1_* amplification |
| p4_fw | pTDH3_G2_fw | CGTGCGAUATAAAAAACACGCTTTTTCAG | ^1^ |
| p4_rv | pTDH3_G2_rv | ATGACAGAUTTTGTTTGTTTATGTGTGTTTATTC | ^1^ |
| p5_fw | A15_fw | CGTGCGAUTTATGGTTTGTGAGGATGTAATC | *CYP716A15* amplification |
| p5_rv | A15_rv | ATGACAGAUTTGTTTTATATTTGTTGTAAAAAGT | *CYP716A15* amplification |
| p6_fw | AaBAS_fw | ATCTGTCAUATGTGGAGATTGAAAATAGCAGA | *AaBAS* amplification |
| p6_rv | AaBAS_rv | CACGCGAUCTAGGTGCCTTTGAGCTGTG | *AaBAS* amplification |
| p7_fw | MTR1_fw | ATCTGTCAUATGACTTCCTCCAATAGTGATCTAGTGCG | *MTR1* amplification |
| p7_rv | MTR1_rv | CACGCGAUTCACCAAACGTCTCTCAAGTACCTAC | Ampliﬁcation of MTR |
| p8_fw | PgDDS_fw | AGTGCAGGUATGTGGAAGCTGAAGGTTGCTCAAGGAAATGACCCTTACTTG | PgDDS amplification |
| p8_rv | PgDDS_rv | CGTGCGAUTTATATTTTAAGCTGCTGGTGTTTAGGC | *PgDDS* amplification |
| p9_fw | A47_fw | AGTGCAGGUATGGCAGCCGCTATGGTTTT | *CYP716A47* amplification |
| p9_rv | A47_rv | CGTGCGAUTCAGTTGTGTGGGTGTAAGTGGAT | *CYP716A47* amplification |
| p10_fw | hphMX-ERG7mut_fw | CTTATTCCCTATTAAGGCATTAGGTATGTACAGCAGGGCATATGAAACACTACGCTTGGAGCAGGTGCTGGTGCTGGTGCTGGAGCAATTCTGTCTAAAGGTGAAGAATTATTCACTGGTGTTGTCCCAATTTTGGTTGAATTAGGACATGGAGGCCCAGAATACCCTCCT | *ERG7* mutation |
| p10_rv | hphMX_ERG7mut_rv | TGCACTAGTTTCTAATTGTTGCAGCCTCTAACAACACTTATAAATAAAACCTTCGAGCGTCCCAAAACCTTCTCAAG | *ERG7* mutation |

Table S4. Triterpenoid production in engineered yeast (combinatorial biosynthesis)

| **Volumetric titer** |  |  | |  | |  | |  | |  | |  | |
| --- | --- | --- | --- | --- | --- | --- | --- | --- | --- | --- | --- | --- | --- |
|  |  | **CDW (g/L)** | | **Lanosterol (mg/L)** | | **Ergosterol (mg/L)** | | **Lupeol (mg/L)** | | **beta-Amyrin (mg/L)** | | **Squalene (mg/L)** | |
| **Strain** | Gene | mean | SD | mean | SD | mean | SD | mean | SD | mean | SD | mean | SD |
| CEN.PK2U K2.3 pTT1-OEW | *OEW* | 12.77 | 0.54 | 4.00 | 0.49 | 57.02 | 0.41 | 87.80 | 4.13 | 0 | 0 | 578.55 | 17.76 |
| CENPK2U K2.3 pTT1-GuLUP1 | *GuLUP1* | 12.50 | 0.04 | 5.25 | 0.15 | 56.24 | 0.63 | 89.36 | 1.59 | 0 | 0 | 629.20 | 54.17 |
| CEN.PK2U K2.3 pTT1-BPW | *BPW* | 12.13 | 0.05 | 18.33 | 0.36 | 52.00 | 1.26 | 0.00 | 0.00 | 0 | 0 | 571.38 | 3.19 |
| CEN.PK2U K2.3 pTT1-OSC3 | *OSC3* | 11.71 | 0.12 | 17.86 | 0.59 | 52.72 | 1.95 | 0.00 | 0.00 | 0 | 0 | 607.57 | 32.99 |
| CEN.PK2U K2.3 pTT1-AtLUP1 | *AtLUP1* | 12.15 | 0.11 | 4.55 | 0.10 | 54.02 | 1.07 | 60.21 | 0.21 | 2.87 | 0.07 | 614.29 | 5.73 |
| CEN.PK2U K2.3 pTT1-RcLUS1 | *RcLUS1* | 12.16 | 0.11 | 4.96 | 0.28 | 53.50 | 2.38 | 88.10 | 2.06 | 0 | 0 | 687.73 | 71.14 |
| **Specific titer** |  |  |  |  |  |  |  |  |  |  |  |  |  |
|  |  | **CDW (g/L)** | | **Lanosterol (mg/g)** | | **Ergosterol (mg/g)** | | **Lupeol (mg/g)** | | **beta-Amyrin (mg/g)** | | **Squalene (mg/g)** | |
| **Strain** | Gene | mean | SD | mean | SD | mean | SD | mean | SD | mean | SD | mean | SD |
| CEN.PK2U K2.3 pTT1-OEW | *OEW* | 12.77 | 0.54 | 0.314 | 0.052 | 4.467 | 0.158 | 6.873 | 0.031 | 0 | 0 | 45.362 | 3.318 |
| CENPK2U K2.3 pTT1-GuLUP1 | *GuLUP1* | 12.50 | 0.04 | 0.420 | 0.013 | 4.500 | 0.067 | 7.150 | 0.152 | 0 | 0 | 50.349 | 4.512 |
| CEN.PK2U K2.3 pTT1-BPW | *BPW* | 12.13 | 0.05 | 1.512 | 0.036 | 4.288 | 0.123 | 0.000 | 0.000 | 0 | 0 | 47.115 | 0.469 |
| CEN.PK2U K2.3 pTT1-OSC3 | *OSC3* | 11.71 | 0.12 | 1.525 | 0.066 | 4.502 | 0.212 | 0.000 | 0.000 | 0 | 0 | 51.885 | 3.341 |
| CEN.PK2U K2.3 pTT1-AtLUP1 | *AtLUP1* | 12.15 | 0.11 | 0.374 | 0.011 | 4.447 | 0.129 | 4.956 | 0.027 | 0.236 | 0.004 | 50.559 | 0.015 |
| CEN.PK2U K2.3 pTT1-RcLUS1 | *RcLUS1* | 12.16 | 0.11 | 0.408 | 0.019 | 4.400 | 0.156 | 7.246 | 0.105 | 0 | 0 | 56.539 | 5.350 |

Table S5. Synthetic genes used in this study

| **Gene (Abbreviation)** | **Origin** | **Enzyme** | **Gene accession number** | **Protein accession number** | **Manufacturer** | **Ref.** |
| --- | --- | --- | --- | --- | --- | --- |
| *OEW* | *Olea europaea* | Lupeol synthase; monofunctional/ oxidosqualene cyclase | AB025343 | Q9SLW3 (UniProt) | GenScript (New Jersey, USA) | ^6^ |
| *GuLUP1* | *Glycyrrhiza uralensis* | Lupeol synthase; monofunctional/ oxidosqualee cyclase | AB663343 | G9MAK0 (UniProt) | GenScript (New Jersey, USA) | ^7^ |
| *BPW* | *Betula platyphylla (var . japonica)* | Lupeol synthase | AB055511 | Q8W3Z2 (UniProt) | GenScript (New Jersey, USA) | ^7^ |
| *OSC3* | *Lotus japonicus* | Lupeol synthase | AB181245 | BAE53430.1 | GenScript (New Jersey, USA) | ^8^ |
| *RcLUS1* | *Ricinus communis* | Lupeol synthase | NM_001323755.1 | NP_001310684.1  Q2XPU7 (Uniprot) | GenScript (New Jersey, USA) | ^9^ |
| *AtLUP1* | *Arabidopsis thaliana* | Lupeol synthase; multifunctional/ oxidosqualene synthase | NM_179572.2 | NP_849903.1 | GenScript (New Jersey, USA) | ^10^ |
| *PgDDS* | *Panax ginseng* | Dammarenediol II synthase, monofunctional | AB265170 | ACZ71036.1  Q08IT1 (UniProt) | IDT (Leuven, Belgium) | ^11^ |
| *LjCPR2* | *Lotus japonicus* | cytochrome P450 reductase, class II | AB433810 | B5BSX2 (UniProt) | GenScript (New Jersey, USA) | ^12^ |
| *ATR1* | *Arabidopsis thaliana* | cytrochrome P450 reductase, class I | NM_118585.4 | NP_194183.1 | GenScript (New Jersey, USA) | ^13^ |
| *ATR2* | *Arabidopsis thaliana* | cytrochrome P450 reductase, class II | NM_119167.4 | Q9SUM3 (UniProt) | GeneArt (Thermo Fisher Scientific Inc., USA) | ^14^ |
| *CrCPR2* | *Catharanthus roseus* | cytrochrome P450 reductase | X69791 | **Q05001 (Uniprot)** | GenScript (New Jersey, USA) | ^15^ |
| *MTR1*(MTR_3g100160) | *Medicago truncatula* | cytrochrome P450 reductase, class I | XM_003602850 | XP_003602898.1, G7J4L3 (UniProt) | GenScript (New Jersey, USA), IDT (Leuven, Belgium) | ^16^ |
| *NCP1* | *Saccharomyces cerevisiae* | cytrochrome P450 reductase | NP_011908.1 | P16603 (UniProt) | GenScript (New Jersey, USA) | ^17^ |
| CYP716A15 | *Vitis vinifera* | C28 oxidase | AB619802 | BAJ84106.1 (Entrez)  F1T282 (UniProt) | GenScript (New Jersey, USA) | ^7^ |
| CYP716A17 | *Vitis vinifera* | C28 oxidase | AB619803 | BAJ84107.1 (Entrez)  F1T283 (UniProt) | GenScript (New Jersey, USA) | ^7^ |
| CYP716A12 | *Medicago truncatula* | C28 oxidase | DQ335781 | ABC59076.1 (Entrez)  Q2MJ20 (UniProt) | GenScript (New Jersey, USA) | ^7^ |
| CYP716AL1 | *Catharanthus roseus* | C28 oxidase | JN565975.1 | AEX07772.1 (Entrez)  I1TEM3 (UniProt) | GenScript (New Jersey, USA) | ^18^ |
|  |  |  |  |  | GeneArt (Thermo Fisher Scientific Inc., USA) | ^1^ |
| CYP716A9 | *Populus trichocarpa* | C28 oxidase | XM_002331391 |  | GenScript (New Jersey, USA) | ^19^ |
| CYP716B2-like | *Glycine max* | C28 oxidase |  | Uniprot: Q50EK0 | GenScript (New Jersey, USA) | BioProject: [PRJNA48389](http://www.ncbi.nlm.nih.gov/bioproject/PRJNA48389) |
| CYP716A41 | *Bupleurum chinense* | C28 oxidase | JF803813 |  | GenScript (New Jersey, USA) | ^20^ |
| CYP716B1-like | *Cucumis sativus* | C-28 oxidase | XM_004139039 |  | GenScript (New Jersey, USA) | BioProject: [PRJNA182750](http://www.ncbi.nlm.nih.gov/bioproject/PRJNA182750) |
| CYP716A47 |  | multifunctional oxidase, acts on C-12 of dammarenediol-II | JN604536.1 | AEY75212.1 | Integrated DNA Technology | ^21^ |

### Characterization of suitable promoters for controlling Erg7 activity

In a previous publication on TIPI (TEV protease induced protein instability), the very tightly regulated promoter of serine/threonine protein kinase (IME2), which is involved in the activation of meiosis, was used for expression control.^22^ Thus, to use TEV protease in this project, a promoter had to be chosen that very tightly controls gene expression and responds well to an external factor for induction to ensure effective protein degradation. Possible promoters include the copper-inducible promoter p*_CUP1_*, or glucose regulated promoters, such as P*_HXT7_* (*HXT7* encodes a high affinity glucose transporter) and P*_ADH2_* (*ADH2* encodes glucose repressed alcohol dehydrogenase II), both of which are repressed during growth in glucose rich medium, as is the case during the first exponential growth phase.

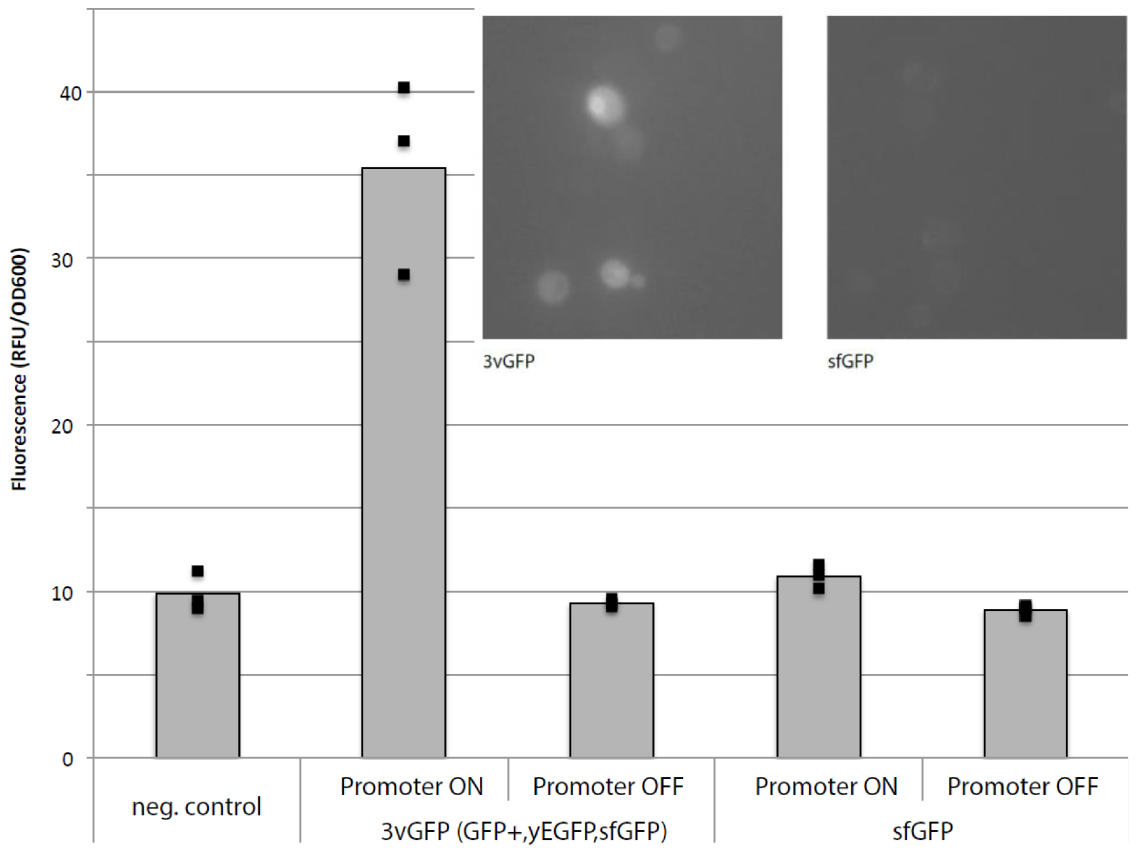

Figure S1. Comparison of the fluorescence of 3vGFP and sfGFP with the autofluorescence of the reference strain *S. cerevisiae* CEN.PK.

Promoters were characterized by using them to control expression of a green fluorescent protein (GFP). Detection of low base line activity was enabled by using a triple tandem protein 3vGFP, which is a fusion protein of three GFP variants (yeGFP-GFP+-sfGFP). ^23^ The P*_CUP1_*-3vGFP construct showed significantly increased fluorescence compared to the autofluorescence of the wild-type control (Fig. S1) as previously described. ^23^

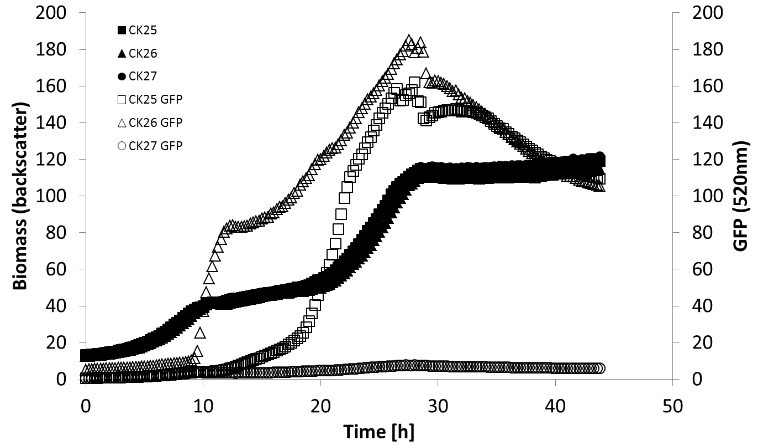

Figure S2. Course of GFP signal (520nm, empty symbols) and biomass (backscatter, filled symbols) of strains CK25 (P*_ADH2_*-3vGFP, squares), CK26 (P*_HXT7_*-3vGFP, triangles) and CK27 (P*_ERG7_*-3vGFP, circles) during cultivation in Delft medium with 2% glucose in the mini-bioreactor system Biolector at 30°C and 1000 rpm.

Selected promoters and the native *ERG7* promoter were cloned by USER cloning upstream of 3vGFP into the integrative EASY clone vector pCFB258 and then stably integrated into the genome of the parent strain CFB1010 at Easyclone integration site 3 on chromosome 10 (X-3).^3^ Strains CK25 (P*_ADH2_*), CK26 (P*_HXT7_*), and CK27 (p*_ERG7_*) were cultured at microtiter scale in the mini-bioreactor system Biolector (m2p labs, Baesweiler, Germany) and data on biomass concentration and GFP signal were recorded at 15 min intervals (Fig. S2).

The strains showed no significant differences with respect to growth, but the course of the GFP signal was different, as expected. For strain CK25 (P*_ADH2_*), the signal was very weak at first, then increased slightly after the end of the glucose phase and showed a steep increase at the beginning of the ethanol phase. The GFP signal of strain CK26 (P*_HXT7_*) started at a slightly elevated level at the beginning and remained relatively constant until just before the end of the glucose phase, when it increased steeply. During the second growth phase, the increase again weakened.

Strains CK24 and CK28 were tested under similar conditions as described above, but here cultivations with and without copper were compared. This comparison showed a significant increase in GFP signal in the cultivations of CK24 in medium containing copper (Fig. S3). For strain CK28, no significant difference was found between the copper concentrations. This could be explained by the signal being too weak to get above the background signal of the medium. To detect finer differences, samples of a similar cultivation were washed and the GFP signal was quantified in a microtiter plate reader (Fig. S3). Here, the uninduced signal (no copper in the medium) was found to be very close to the autofluorescence of the wild-type strain, but the signal induced by copper was significantly increased.

We also tested the copper inducible *CUP1* promoter (P*_CUP1_*). The strains with the tagged Erg7p protein and TEV protease under P*_CUP1_* control were able to accumulate 2,3-oxidosqualene. However, this occurred even without the addition of copper to the medium, suggesting that the control by P*_CUP1_* is not stringent enough. To increase stringency, a fusion promoter consisting of the *CUP1* promoter and part of the *SPO13* (meiotic regulator) promoter was constructed. *SPO13* is highly regulated by two *UME6* (Unscheduled Meiotic gene Expression) binding sites.^23^ By combining the inducible copper promoter and the tightly controlled SPO13, basal expression is greatly reduced, but induction is still ensured by the addition of copper (Fig. S3).

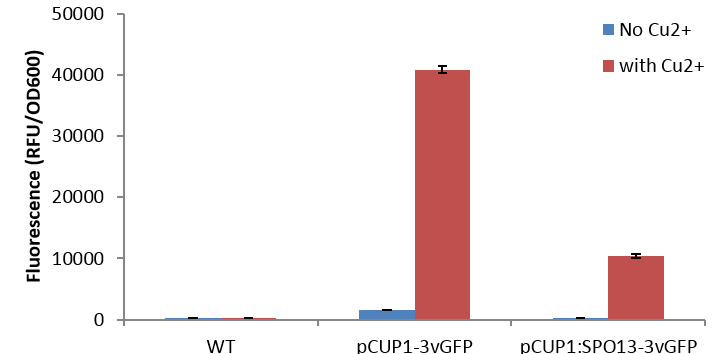

Figure S3. Fluorescence of the wild-type strain, p*_CUP1_*-3vGFP (CK24), and p*_CUP1_*-SPO13-3vGFP strain (CK28) measured after cultivation in medium with (red) and without (blue) copper.

The promoter characterization results show that both glucose-dependent systems (P*_ADH2_*, P*_HXT7_*) have a very good response, but the *HXT7* promoter has an increased basal activity, which could already lead to increased TEV protease expression. The results for the characterization of the CUP1 promoter show that it has too much basal activity and therefore cannot be recommended for the degron strategy. The fusion promoter *CUP1*-*SPO13* showed promising activity behavior, with very weak basal activity and good inducibility.
